# The YTHDC1 glutamate-rich domain docks to the ADAR1 Zα Domain, linking the N^6^-methyladenosine modification of pre-mRNAs to dsRNA editing

**DOI:** 10.64898/2026.07.08.737312

**Authors:** Dmitry Gromak, Alexey K. Shaytan, Alan Herbert, Maria Poptsova

**Affiliations:** HSE University, Moscow, Russia; Lomonosov Moscow State University, Moscow, Russia; InsideOutBio, Charlestown, MA, USA

**Keywords:** ADAR1, YTHDC1, Zα domain, A-to-I RNA editing, m6A epitranscriptomics, protein–protein interaction, AI, protein design, RFdiffusion, molecular dynamics simulation, innate immunity

## Abstract

The p150 isoform of the double-stranded RNA editing enzyme ADAR1 binds Z-DNA and Z-RNA through the conserved winged helix-turn-helix Zα domain. Here, we describe an inverse computational design strategy to map protein interactors of Zα. We used RFdiffusion and ProteinMPNN to generate ∼10,000 synthetic binders optimized for the Zα recognition surface, then used their sequences as structural templates for BLASTp searches against the human proteome. Multi-stage screening of ∼1,200 candidate regions from 298 proteins via ColabFold pDockQ identified 79 candidates for high-resolution AlphaFold3 modeling, which revealed the m6A reader YTHDC1 as the top-ranked interactor. AlphaFold3 predicts that a glutamate-rich poly-E disordered region of YTHDC1 (residues 199–254) docks into the basic recognition pocket of Zα through a charge-complementary mechanism that mimics the phosphate backbone of Z-RNA. Microsecond molecular dynamics simulations confirmed stability of the binary ADAR1p150–YTHDC1 complex, with the Zα–poly-E interface maintaining RMSD below 3 Å throughout. Ternary complex simulations showed that dsRNA acts as a co-anchoring scaffold stabilizing simultaneous engagement of both proteins in a catalytically dormant conformation. YTHDC1 localizes to transcription-associated YT bodies where nascent RNAs undergo m6A modification and negative supercoiling promotes Z-DNA formation, suggesting that YTHDC1 recruits ADAR1p150 to promote editing of intron-containing substrates prior to splicing.

**Short Statement:** ADAR1 can edit RNAs after they are made, changing the message they carry and removing double-stranded RNAs that can activate inflammatory responses. Using AI-driven protein design, we discovered that ADAR1 physically interacts with YTHDC1, a protein that recognizes newly made self-RNAs. The interaction allows ADAR to edit RNAs as they are made. This unexpected connection between two fundamental RNA modification systems opens new avenues for understanding and potentially targeting autoimmune disorders and cancers driven by the mis-editing of RNA transcripts.

## Introduction

Adenosine Deaminase Acting on RNA 1 (ADAR1) is a cornerstone of the human innate immune system, responsible for the hydrolytic deamination of adenosine to inosine (A-to-I) within double-stranded RNA (dsRNA) (1,2). This modification is vital for distinguishing “self” from “non-self” transcripts, preventing endogenous RNAs from triggering an inflammatory response via sensors like MDA5 or PKR (3,4). A defining feature of the interferon-inducible ADAR1 p150 isoform is its N-terminal Zα domain (5-7). While we have a clear structural understanding of how this winged helix-turn-helix motif recognizes left-handed Z-conformations of nucleic acids (8), its potential to act as a scaffold for protein-protein interactions (PPIs) has remained largely unexplored. The biological versatility of ADAR1 suggests it may be recruited to various RNA-processing hubs, yet many of its functional partners remain unknown. We hypothesized that the Zα domain might serve as a conformation-specific recruiter, docking with protein motifs that mimic the geometry or charge of Z-form nucleic acids. In this study, we set out to map the potential Zα interactome using a broad, systematic search. Rather than beginning with a specific list of suspects, we used an “inverse design” strategy: we first used generative AI to design the “perfect” synthetic binders for the Zα surface and then searched the human proteome for natural sequences that matched these structural templates. This pipeline, which integrates RFdiffusion (9), ProteinMPNN (10), AlphaFold3 (11), and long-scale molecular dynamics, successfully identified the m6A reader YTHDC1 as a novel interactor, providing a physical link between two major pathways of the human epitranscriptome.

## Materials and Methods

### *De Novo* Design and BLAST Mapping

The target structure for the Zα domain was derived from the crystal structure of the ADAR1 Zα domain in complex with Z-RNA (PDB ID: 2GXB) (7); chain D (author chain B) was isolated and used as the receptor conformation for all generative design runs. We used RFdiffusion (9,12) to generate 1,200 backbones in four batches. Batches 3 and 4 utilized a custom potential based on the backbone RMSD to the C2′ atoms of a reference dsZ-DNA structure. Batches 2 and 4 incorporated hotspots for residues K169, K170, N173, R174, and Y177. The target sequences were designed with ProteinMPNN (10) (8 per backbone). Initial filtering in ColabFold v1.5 (13) required pLDDT > 80, iPAE < 10 Å, and interface backbone RMSD < 3 Å. Mapping was performed using BLASTp against SWISS-PROT (14) with an e-value cutoff of 1 × 10^-5.

### Structural Refinement and Scoring

Candidate regions were folded with Zα in ColabFold, and the 79 resulting hits were modeled as full-length complexes using the AlphaFold3 server (11). To mitigate the impact of disordered regions on global PAE scores, we used pDockQ pDockQ (15) as our primary differentiation metric. A threshold of ≥ 10 inter-chain contact pairs (Cα-Cα distance < 8 Å) was applied to filter out random alignments.

### Molecular Dynamics Simulations

MD simulations were performed using the Amber24 software package (19). The models were placed in a rectangular box with a minimum distance between model and box set to 12 Å. Amber force field ff19SB with OL3 force field for RNA and an OPC water model were used. Na^+^ and Cl^−^ ions were added to neutralize the system. Production runs were conducted at 300 K (Langevin dynamics for temperature regulation) and 1 bar (Berendsen barostat) with 10 Å non-bonded cutoff radius and a 2 fs time step (dt = 0.002) for 5 × 10^8 steps. Trajectory analysis was performed using the cpptraj module of AmberTools25. Stability was assessed by protein backbone RMSD (calculated relative to the starting structure) and residue-level RMSF(calculated relative to the time-averaged structure over the production trajectory).

## Results

### Generation of ADAR1 Z*α* protein binders

We used RFdiffusion (9,12) to generate a library of potential binders, exploring the target surface typically occupied by the phosphate backbone of Z-RNA. To ensure a thorough sampling of the binding landscape, we produced 1,200 backbones across four batches: (1) unconditional generation; (2) hotspot-conditioned generation targeting key residues like K169, K170, and Y177; (3) generation using a custom potential that favored backbone atom alignment to the C2′ atoms of a reference dsZ-DNA structure; and (4) a hybrid approach combining both hotspots and conformational potentials. Each backbone was assigned eight sequences via ProteinMPNN (10), yielding a starting library of approximately 10,000 designs. These were initially validated by folding them with the Zα domain in ColabFold v1.5 (13). While most designs showed high confidence in local folding (pLDDT > 80) and relative orientation (iPAE < 10 Å), we noticed that many backbones deviated significantly from their designed coordinates during the folding process. We initially applied a strict 2 Å RMSD filter, but this was too restrictive, yielding only five hits. By relaxing the threshold to 3 Å, we isolated 11 high-quality binders that maintained the intended interface geometry and served as our templates for proteome mapping (Figure 2, Supplemental Data 1).

**Figure 1.**
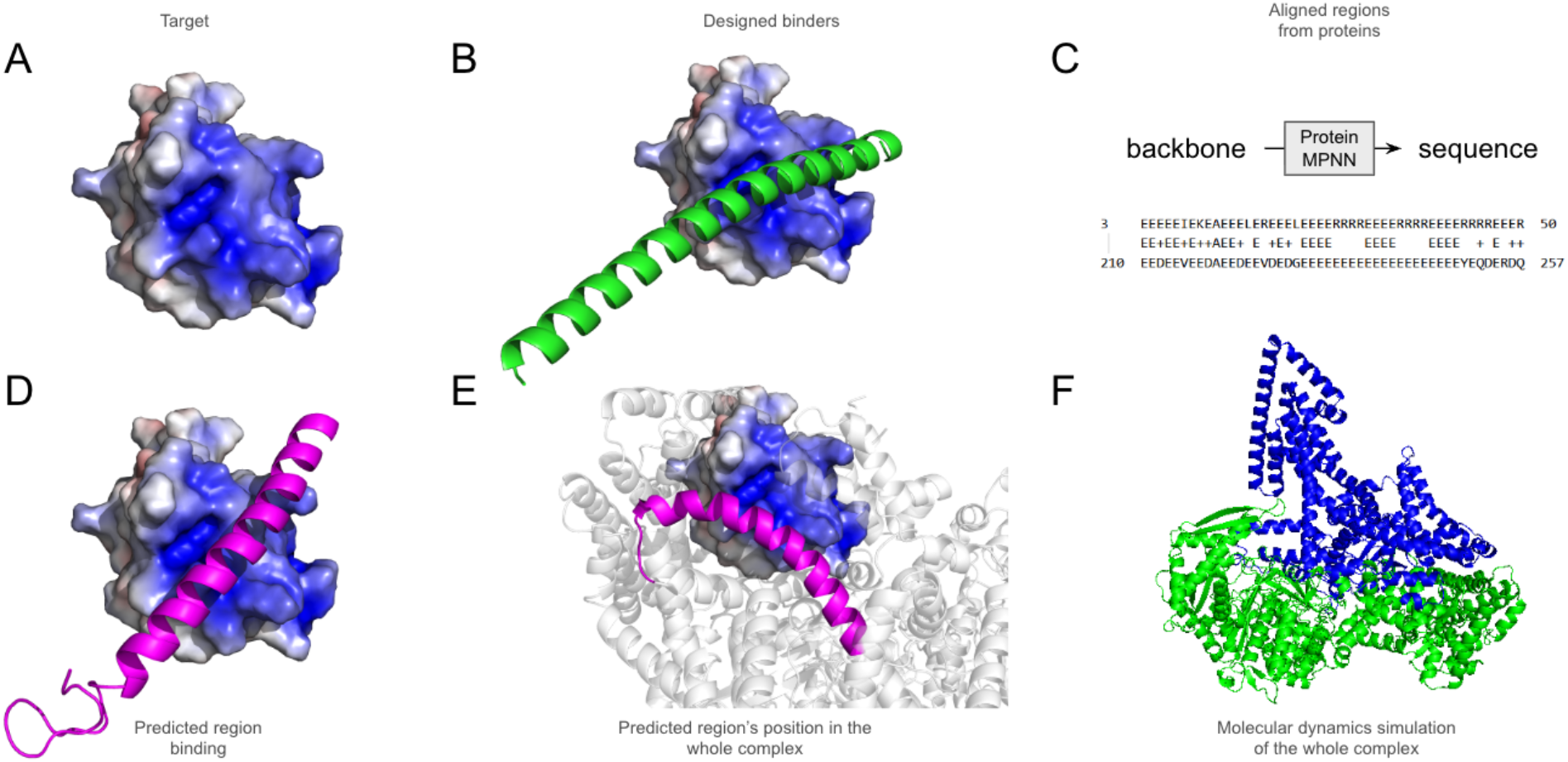
Computational pipeline for identifying Zα-domain protein interactors. A. Electrostatic surface representation of Zα domain target. B. *De novo* binder backbone generated by RFdiffusion. C. Prediction of amnio-acid sequences with protein MPNN and BLASTp alignment of the generated amino acid sequence to YTHDC1 (residues 210–257). D. ADAR1 Zα-domain with the corresponding region from YTHDC1 (shown as magenta on the Zα surface). E. YTHDC1 and Zα-domain of ADAR1 (the candidate binding region of YTHDC1 positioned within the context is highlighted in magenta). **F.** ADAR1-YTHDC1 interactions are confirmed by molecular dynamics simulation of the entire complexes (ADAR1, green; YTHDC1, blue).

**Figure 2.**
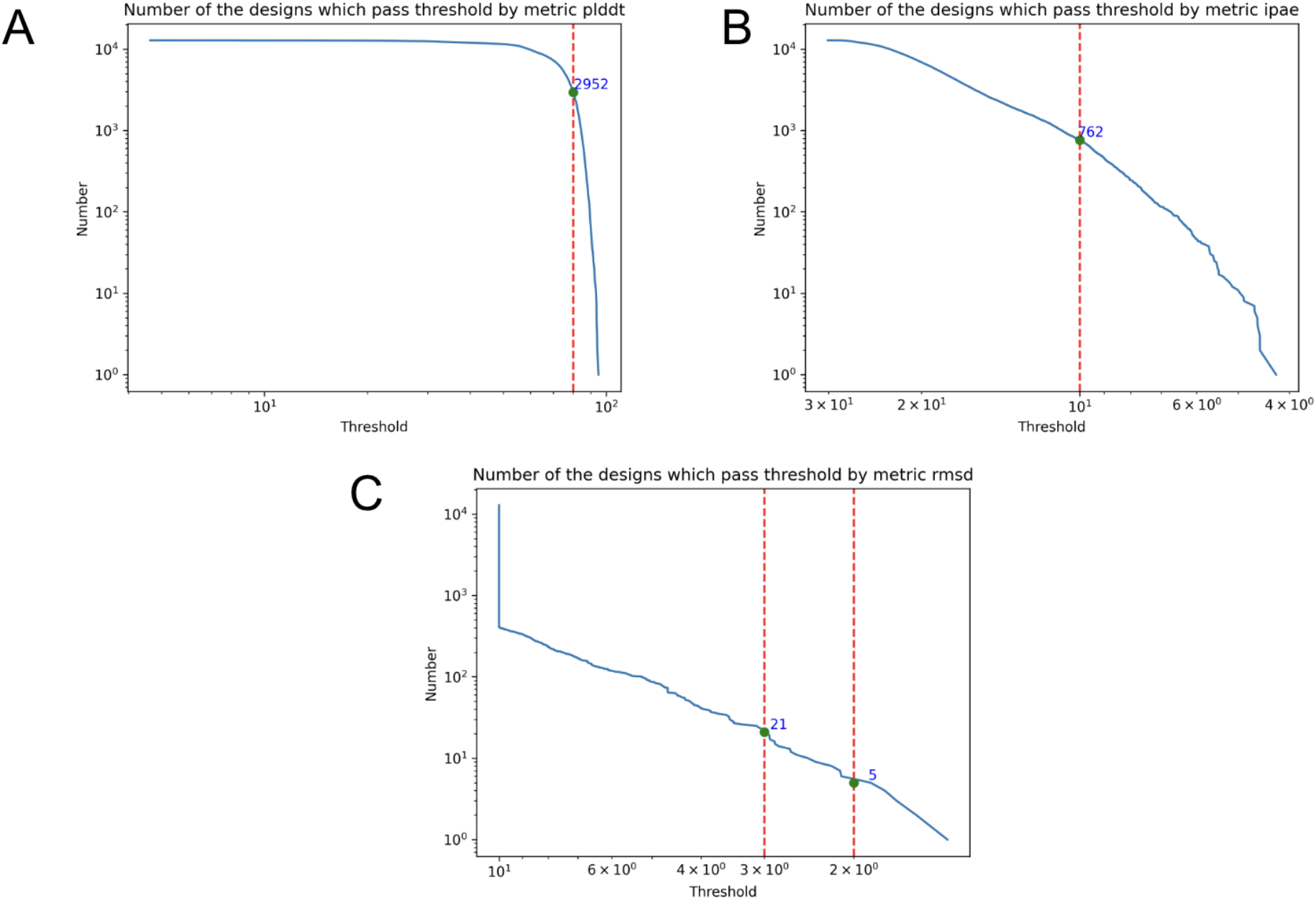
Quality-filter threshold curves for the de novo RFdiffusion binder library (n ≈ 10,000 designs). A. Number of designs passing the pLDDT threshold (dashed line, pLDDT > 80; 2,952 designs at cutoff). B. Number of designs passing the iPAE threshold (dashed line, iPAE < 10 Å; 762 designs at cutoff). C. Number of designs passing the interface backbone RMSD threshold (left dashed line, RMSD < 3 Å, 21 passing designs; right dashed line, RMSD < 2 Å, 5 passing designs).

### Proteome Mapping and Multi-Stage Candidate Screening

To transition from synthetic designs to biological reality, we used the sequences of our 11 lead binders as queries for BLASTp searches against the SWISS-PROT database (14). By setting an e-value threshold of 1 × 10^-5, we identified approximately 1,200 regions across 298 proteins that shared significant similarity with our designed binders. These regions were extracted and co-folded with the Zα domain in a high-throughput screen. To identify the most promising hits for full-length modeling, we evaluated four metrics: pDockQ (15), pDockQ2 (16), ipTM (17), and ipSAE (18). As thresholds for these metrics became more stringent, the pool of successful candidates narrowed from hundreds down to dozens (Figure 3). We selected the 79 proteins that consistently passed the lower thresholds of all four metrics to move forward into high-resolution modeling.

**Figure 3.**
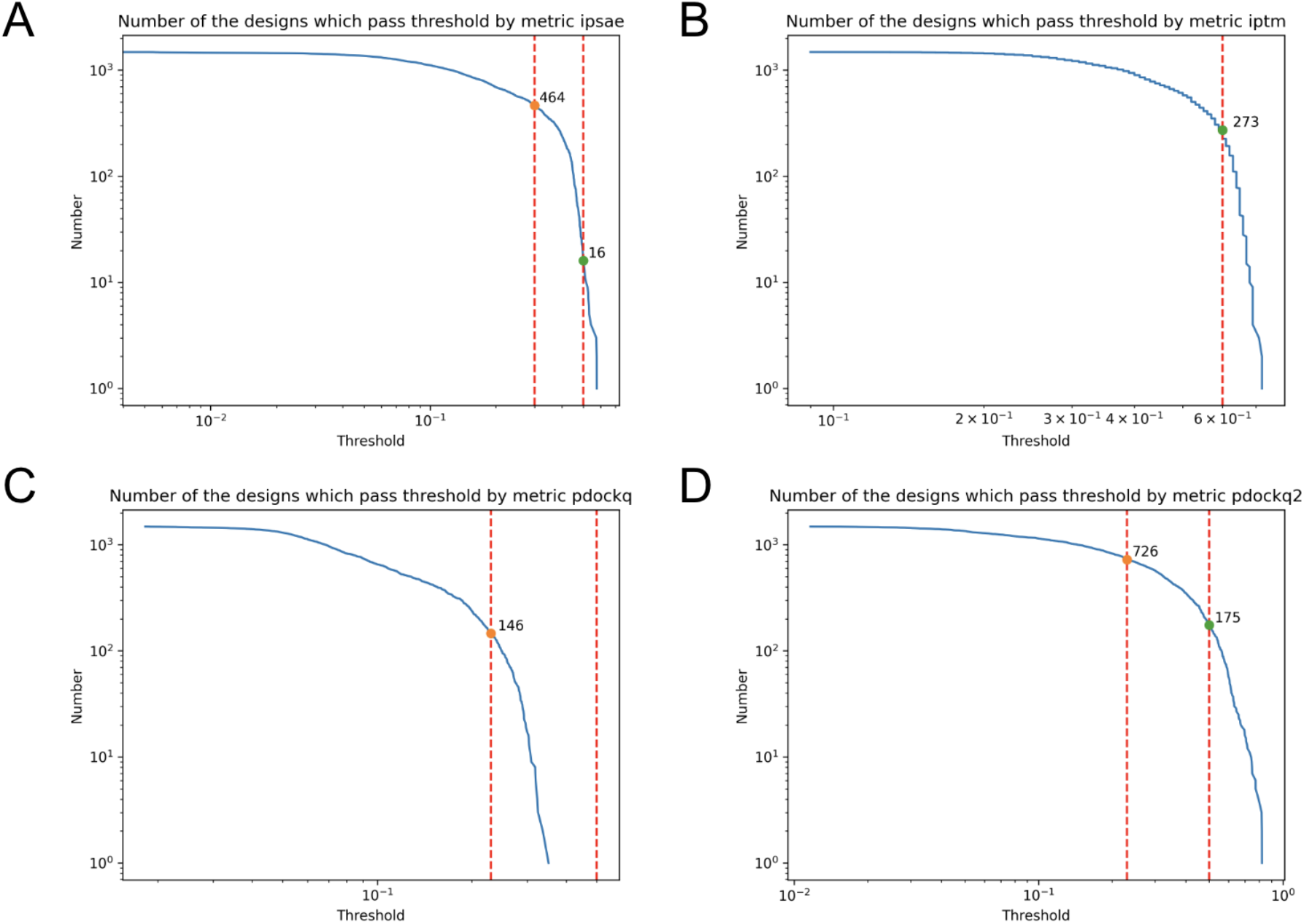
Multi-metric threshold analysis of ∼1,200 candidate regions from 298 proteins identified by BLASTp homology search against SWISS-PROT. A. Number of candidate regions passing the ipSAE threshold (orange marker, lower cutoff, 464 regions; green marker, upper cutoff, 16 regions). B. Number of candidate regions passing the ipTM threshold (lower cutoff, 273 regions). C. Number of candidate regions passing the pDockQ threshold (lower cutoff, 146 regions). D. Number of candidate regions passing the pDockQ2 threshold (lower cutoff, 726 regions; upper cutoff, 175 regions). Dashed red vertical lines indicate applied threshold boundaries; 79 proteins passing lower thresholds across all four metrics advanced to full-length AlphaFold3 modeling.

### AlphaFold3-predicted ADAR1-YTHDC1 complex

We used the AlphaFold3 (AF3) server (11) to model these 79 candidates in their full-length forms with ADAR1 p150 isoform. We found that standard global confidence scores like ipTM and ipSAE were often quite low, even for models that looked visually compelling. We think that this was a result of the extensive intrinsically disordered regions (IDRs) in both ADAR1 and its potential partners; the high “aligned error” (PAE) in these flexible tails tends to wash out the signal from localized binding interfaces. To bypass this limitation, we prioritized the pDockQ metric, which focuses specifically on the density and confidence of contacts at the interface rather than the entire protein chain. Sorting the results by pDockQ provided a clear differentiation of the interactome (Figure 4). We excluded any complexes where the Zα domain formed fewer than 10 contact pairs with the target region, leaving 23 high-confidence candidates. Among these, YTHDC1 stood out as a particularly striking hit. The model predicted that an acidic “poly-E” motif in YTHDC1 (residues 199–254) docks directly into the basic recognition pocket of Zα, essentially acting as a molecular mimic of the negatively charged Z-RNA backbone (Figure 5).

**Figure 4.**
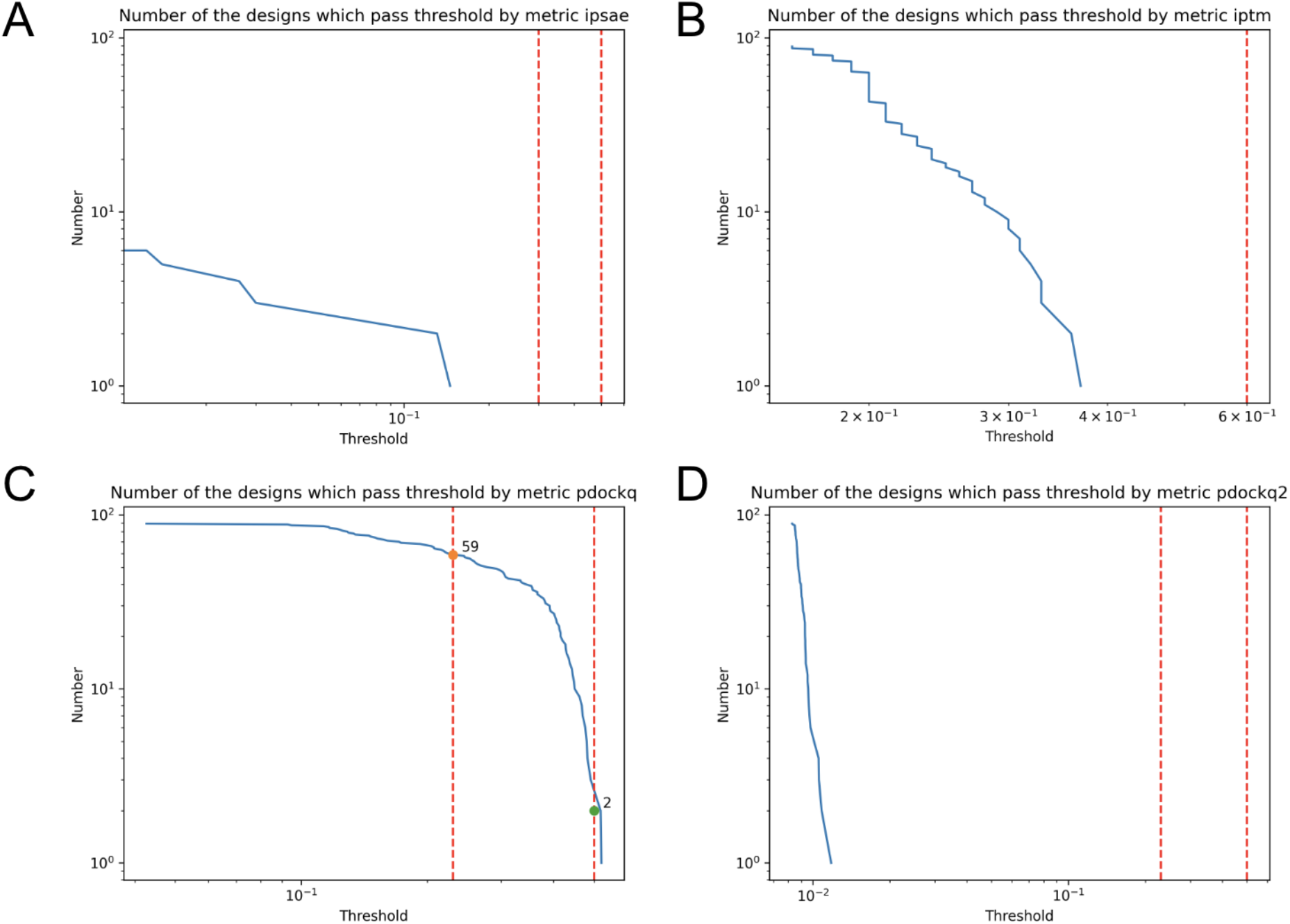
Metric threshold analysis of 79 full-length ADAR1 p150 complexes modeled by AlphaFold3. A. Number of full-length complexes passing the ipSAE threshold. B. Number of full-length complexes passing the ipTM threshold. C. Number of full-length complexes passing the pDockQ threshold (orange marker, lower cutoff, 59 complexes; green marker, upper cutoff, 2 complexes). D. Number of full-length complexes passing the pDockQ2 threshold. Dashed red vertical lines indicate applied threshold boundaries; 23 complexes with ≥ 10 Cα–Cα inter-chain contact pairs passing the upper pDockQ cutoff as high-confidence candidates.

**Figure 5.**
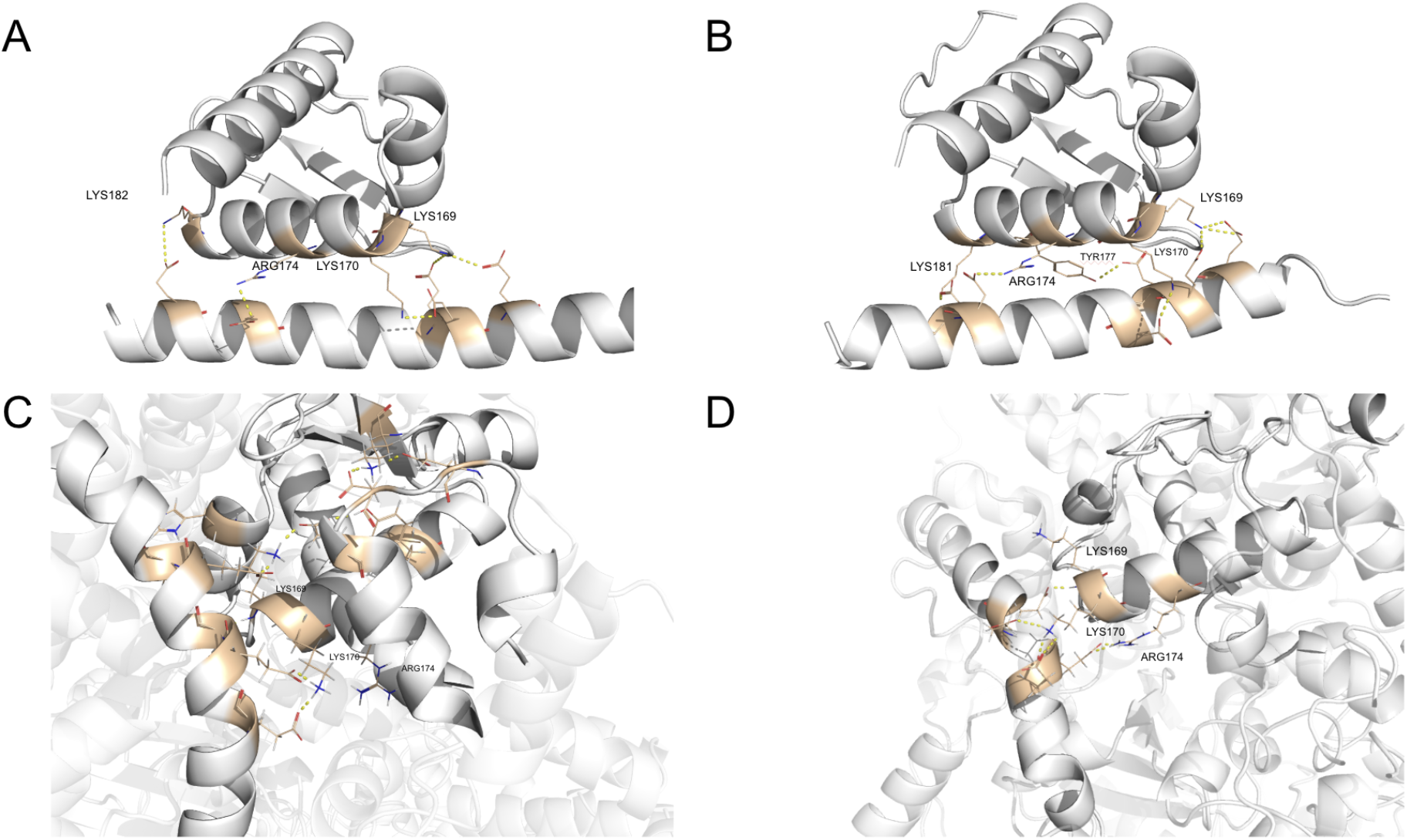
Zα-domain interface residue contacts at four stages of the computational pipeline. A. RFdiffusion-designed binder co-folded with the Zα domain, with hydrogen bonds (yellow dashes) to key basic residues LYS169, LYS170, ARG174, and LYS182. B. Isolated YTHDC1 poly-E region (residues 199–254) co-folded with Zα, with contacts to LYS169, LYS170, ARG174, TYR177, and LYS181. C. Poly-E motif within the full AlphaFold3-predicted ADAR1p150–YTHDC1 complex, with the same charge-complementary interface to LYS169, LYS170, and ARG174. D. Zα–poly-E interface after 1 µs of explicit-solvent MD simulation, with retained contacts to LYS169, LYS170, and ARG174.

### Atomic Stability of ADAR1-YTHDC1 complex confirmed by MDS

To confirm that the AlphaFold 3 predicted ADAR1p150–YTHDC1 complex is physically stable, we performed a 1 µs molecular dynamics (MD) simulation using Amber (19). The assembly reached structural equilibrium within 200 ns, maintaining a stable backbone RMSD of ∼8 Å. For a complex of this size with multiple disordered linkers, this value indicates a well-preserved global structure. Most importantly, the Zα -polyE interface was remarkably rigid, with an RMSD staying below 3 Å throughout the entire trajectory (Figure 6).

**Figure 6.**
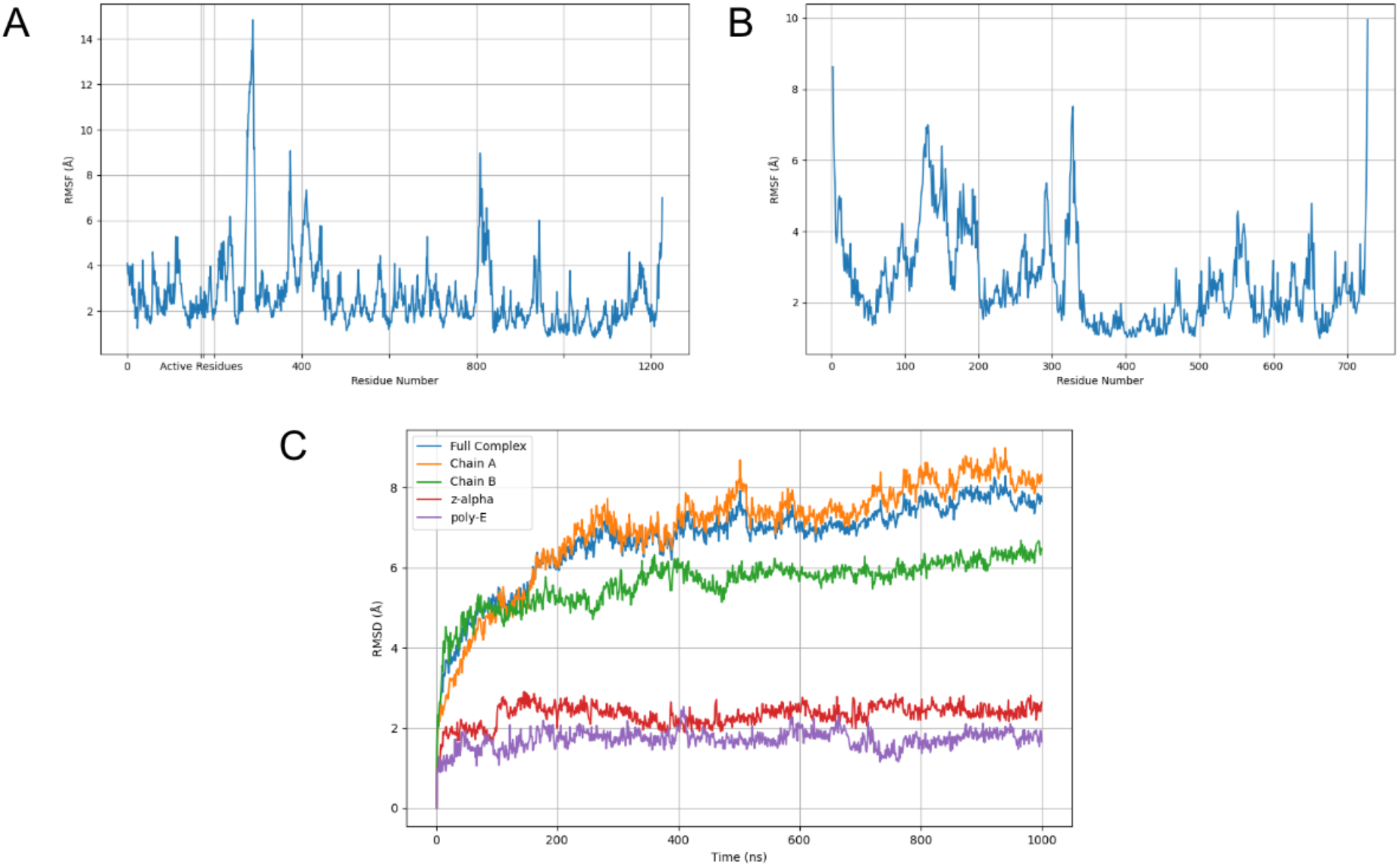
Structural stability of the binary ADAR1p150–YTHDC1 complex over 1 µs of MD simulation. A. Per-residue backbone RMSF profile across the ADAR1 with LYS169, LYS170 and ARG174 labeled as active residues. B. Per-residue backbone RMSF profile across the YTHDC1. C. Backbone RMSD over simulation time for the full complex (blue), ADAR1p150/chain A (orange), YTHDC1/chain B (green), the Zα domain (red), and the poly-E motif (purple), with equilibrium reached by ∼200 ns and interface RMSD below 3 Å throughout.

### RNA-Mediated Assembly of ADAR1–YTHDC1

Given ADAR1’s primary role in RNA binding, we also investigated whether this interaction could persist in the presence of RNA. Using an RNA scaffold derived from the 9B89 PDB structure [https://doi.org/10.2210/pdb9B89/pdb], we modeled a ternary complex. The resulting 1 µs MD simulation showed that the assembly is highly stable, with the RNA acting as a central axis that anchors both the ADAR1 and YTHDC1 molecules (Figure 7). This suggests that YTHDC1 can engage ADAR1 at transcriptionally active sites without competing with Z-RNA, potentially even being stabilized by it.

**Figure 7.**
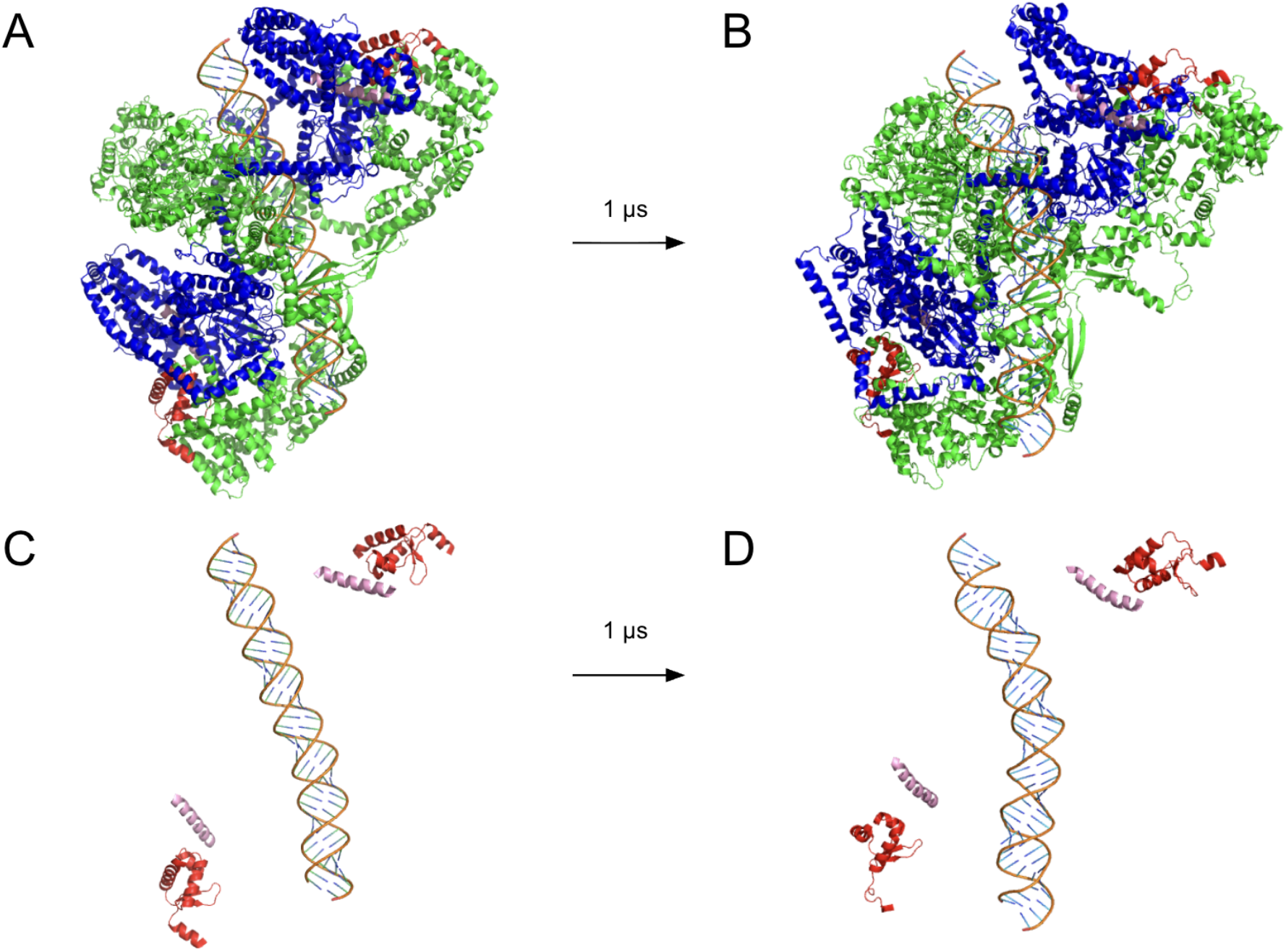
Structural snapshots of the ternary ADAR1p150(dimer)–YTHDC1(dimer)–dsA-RNA complex before and after 1 µs of MD simulation. A. AF3-predicted starting conformation of the full heterotetramer (ADAR1 protomers, green; YTHDC1 protomers, blue; dsA-RNA scaffold, orange). B. Conformation of the full heterotetramer after 1 µs, with preserved global domain arrangement. C. Zoomed view of the isolated Zα–dsA-RNA–poly-E sub-complex at the start of simulation. D. Zoomed view of the sub-complex after 1 µs, with both proteins co-anchored to the Z-RNA helical axis.

To quantify the structural integrity of the full heterotetramer, we analyzed chain-level and interface-level Cα RMSD trajectories throughout the 1 µs simulation (Figure 8). All four protein chains reached equilibrium by approximately 200 ns, plateauing below ∼6 Å – a level consistent with a stable, non-dissociating assembly of this size and IDR content. Strikingly, the interface-level analysis revealed that all four Zα–poly-E contacts – one from each ADAR1 protomer paired with one from each YTHDC1 protomer – remained below ∼2.5 Å RMSD throughout the entire trajectory. This symmetric stability across both protomer pairs demonstrates that the charge-complementary interaction is not an artifact of a single interface but is a robust, geometrically equivalent feature of the full dimeric assembly. Furthermore, the observation that the ADAR1 deaminase domain did not translocate toward the RNA substrate during the simulation suggests that the ternary complex adopts a catalytically dormant configuration, in which Z-RNA serves as a structural anchor rather than a direct editing substrate. Together, these data support a model in which Z-RNA nucleation at transcriptionally active loci drives symmetric co-recruitment of ADAR1 and YTHDC1 dimers, forming a stable regulatory platform that physically links A-to-I editing and m6A readout on the same RNA scaffold.

**Figure 8.**
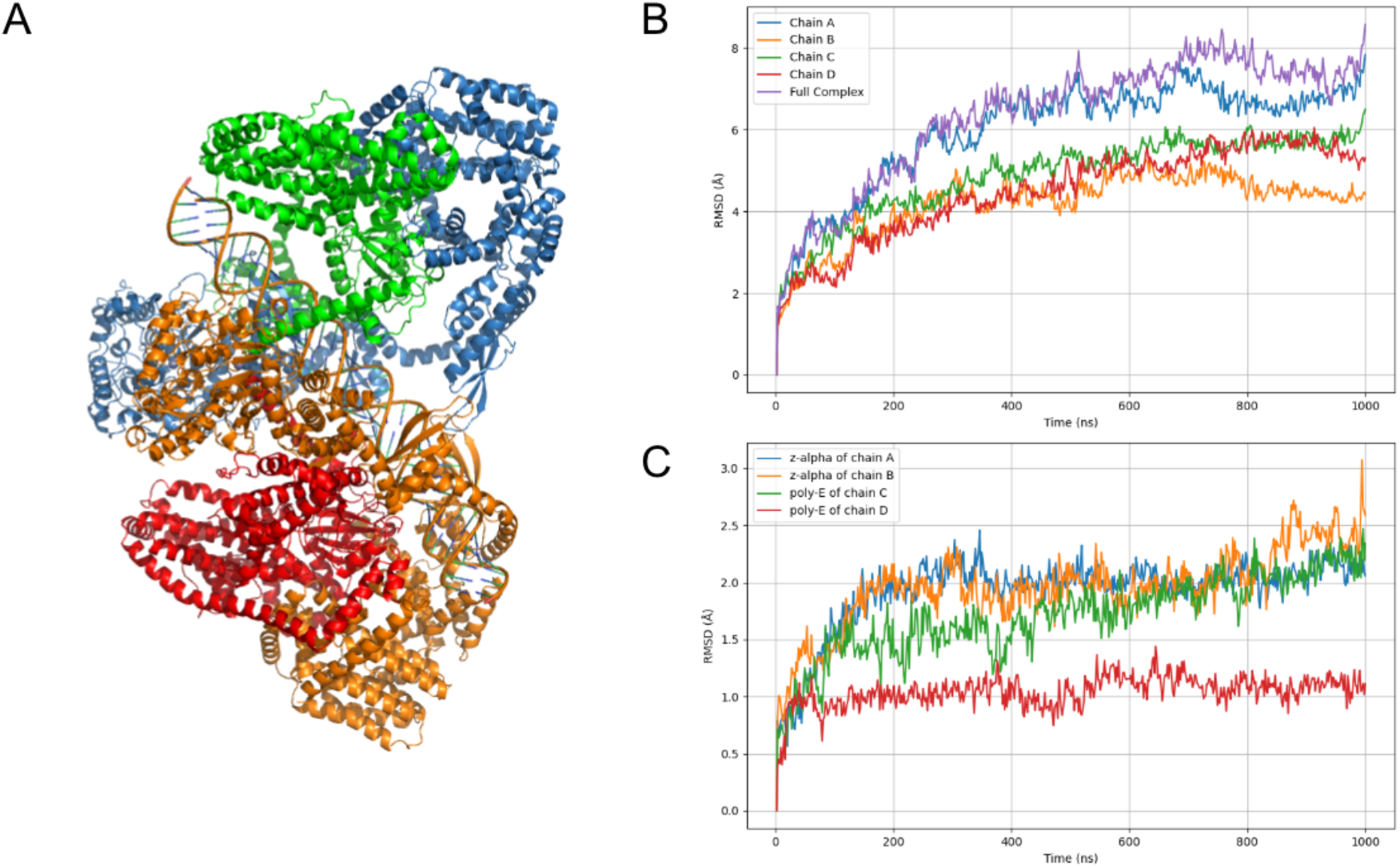
Backbone Cα RMSD over the 1 µs MD trajectory of the ternary ADAR1p150(dimer)– YTHDC1(dimer)–dsA-RNA complex. A. Representative snapshot of the ADAR1p150–YTHDC1– dsZ-RNA ternary complex, colored by chain, matching the color scheme in (B) and (C). B. Chain-level RMSD for each of the four protein chains (A–D) and the full assembly. C. Interface-region RMSD for the Zα domains of both ADAR1 protomers (chains A and B) and the poly-E motifs of both YTHDC1 protomers (chains C and D), with all four interface regions below ∼2.5 Å throughout the trajectory.

## Discussion

The innate immunity sensor ADAR1 and the m6A reader YTHDC1 have each been studied intensively, yet their potential functional relationship has remained unexplored. Here, we leveraged an inverse design strategy – generating synthetic binders for the ADAR1 Zα surface with RFdiffusion and ProteinMPNN, and then mapping their sequences back onto the human proteome – to identify YTHDC1 as a novel interactor. The central finding is that a glutamate-rich (poly-E) intrinsically disordered region of YTHDC1 (residues 199–254) docks into the basic recognition pocket of the Zα domain through a charge-complementary mechanism that structurally mimics the negatively charged backbone of Z-form RNA.

The Zα domain is canonically understood as a structural reader of left-handed Z-RNA or Z-DNA, engaging the nucleic acid phosphate backbone through a conserved set of basic residues (K169, K170, R174, Y177, K181, K182). Strikingly, the AlphaFold3 models and the 1 µs molecular dynamics simulations show that the poly-E helix of YTHDC1 maintains sustained contacts with the same basic patch while preserving the overall geometry of the Zα recognition surface. This supports the hypothesis that the Zα domain can function as a dual-purpose scaffold: a nucleic acid reader in the canonical sense, and a protein interaction hub when the polyanionic charge of Z-RNA is mimicked by an intrinsically disordered protein segment.

Our results suggest that the poly-E motif of YTHDC1 is a high-affinity ligand for Zα. YTHDC1 (also known as Yt521-B) binds to the N^6^-methyl-adenosine (m6A) RNA mark that is generated co-transcriptionally. It is the only m6A reader that is also nuclear. The other family members (YTHD-C2, YTHDF1, YTHDF2, YTHDF3) lack the polyE motif. In the nucleus, YTHD-C1 forms what are called YT-bodies during the S-phase of cell division, when transcription restarts, and retrotransposition of retroelements is observed (20). Formation of YT-bodies depends on transcript elongation, as it is inhibited by actinomycin-D but not by α-amanitin or 5,6-dichloro-1-β- ribofuranosyl-benzimidazole (DRB) (21). Potentially, YTHDC1 provides another mechanism for localizing Zα-containing proteins like ADAR to nascent transcripts, enabling editing of intron-containing double-stranded RNA substrates prior to the removal of introns by splicing. The YT bodies are adjacent to nuclear speckles where the m6A-binding activity of YTHDC1 localizes the splicing factor SRSF3, and excludes SRSF10 (22). This arrangement of YT-bodies and splicing speckles suggests a production line through which mRNA is assembled, with editing and splicing physically separated. The design also allows intron-directed editing within YT-bodies of splice sites and small RNA precursors, such as pri-miRNAs (23). Mutations in the Zα domain that cause Aicardi–Goutières syndrome (AGS) (24,25) would be predicted to disrupt both localization of ADAR p150 to YT bodies, leading to mis-processing and mis-splicing of immune active transcripts and miRNAs (26).

The concept of a protein acting as a nucleic acid mimic is well established in translation and antiviral biology. Interestingly, a previous biochemical study using Zα as a bait identified the C-terminal D/E-rich domain of Methyl-CpG-binding domain protein 3 (MBD3) as a high-affinity ligand that bound Zα in vitro with nM affinity (27). In a competition assay, Zα could outcompete the MDB3 domain for binding to a Z-DNA-forming ligand, but not vice versa. This outcome likely reflects a slower off-rate for Zα compared to the MDB3 domain. MDB3 is part of the Nucleosome Remodeling and Deacetylase (NuRD) complex that remodels chromatin to regulate gene expression. Both nucleosome displacement and the negative supercoiling generated by transcription can induce the transition from B-DNA to Z-DNA. In this scenario, the interaction of the D/E-rich MDB3 domain with Z-DNA binding proteins occurs at sites of Z-DNA formation. In the case of Zα, this may help localize the p150 isoform of ADAR to sites of nascent transcript formation, while in other cases, it may help reinitiate transcription by binding Transcription Factor E (28,29). Interestingly, the experimentally verified interaction has a pdockq score of 0.32, ranking it twentieth on the list of hits given in Supplemental Data 1, below YTDC1

A key methodological contribution of this work is demonstrating that inverse computational design can effectively guide proteome-wide interactome discovery. Standard approaches to identifying ADAR1 partners rely on affinity purification coupled to mass spectrometry or yeast two-hybrid screens, which are inherently limited by expression context, affinity thresholds, and indirect interactions. By generating structurally optimal synthetic binders first, we created a biophysically defined search template that encodes the precise interface geometry and charge topology of the Zα surface. This approach bypassed the noise inherent in sequence-only homology searches and focused attention on proteins with the correct three-dimensional complementarity. The multi-stage screening pipeline – filtering approximately 10,000 designs down to 11 templates, then 1,200 candidate regions down to 79 full-length models – demonstrated that incorporating multiple orthogonal confidence metrics (pDockQ, pDockQ2, ipTM, ipSAE) is essential for controlling false positives, particularly given the prevalence of intrinsically disordered regions in both ADAR1 and its binding partners.

The challenge posed by disordered regions deserves specific attention. Both ADAR1p150 and YTHDC1 contain extensive IDRs that dramatically suppress global PAE-based metrics, such as ipTM and ipSAE, even when the folded interface is well predicted. Our observation that pDockQ, which evaluates contact density at the interface rather than global chain alignment, was the only metric capable of reliably discriminating candidates in full-length AF3 models has practical implications for future computational interactome studies involving IDR-containing proteins. We suggest that interface-local scoring metrics should be prioritized over global confidence scores in such contexts, and that contact-pair filtering (here, ≥10 Cα–Cα pairs within 8 Å) provides a useful secondary filter against spurious high-pDockQ predictions arising from random chain proximity.

The molecular dynamics simulations provide strong evidence that the predicted interaction is not an artifact of static structure prediction. In the binary complex, the Zα–poly-E interface maintained an RMSD below 3 Å over the entire 1 µs trajectory, despite the full complex fluctuating at ∼8 Å –a level of interface rigidity that far exceeds the global structural variability and is indicative of a genuine, thermodynamically stable protein–protein interaction. The ternary simulation with a dsRNA scaffold is more consistent with competitive displacement than with co-occupancy at the Zα surface. The overall assembly remained globally stable – the RNA scaffold was retained through contacts with the dsRBDs – yet the Zα–poly-E interface was maintained throughout the trajectory rather than being dissolved by the competing RNA. This suggests that YTHDC1’s poly-E motif outcompetes Z-RNA for occupancy of the Zα basic recognition pocket, retaining the RNA within the complex through dsRBD contacts alone. The deaminase disengagement observed below is a structural consequence of this displacement: without the Zα–RNA anchor, the dsRNA cannot adopt the geometry required for catalytic engagement of the deaminase. Notably, the deaminase domain of ADAR1 did not adopt an editing-competent conformation during the simulation, suggesting the ternary complex may represent a catalytically dormant, RNA-anchored state rather than an active editing complex. This raises the intriguing possibility that YTHDC1 binding modulates ADAR1 editing activity, perhaps by restricting access of the dsRNA substrate to the deaminase active site or by sequestering ADAR1 at specific chromatin-associated RNA substrates pending a conformational trigger.

Several important questions remain to be addressed experimentally. First, the interaction should be validated biochemically, for example by co-immunoprecipitation, surface plasmon resonance, or isothermal titration calorimetry using the isolated Zα domain and the YTHDC1 poly-E region. The charge-complementary nature of the interface makes it amenable to mutagenesis validation: substitution of the key basic residues on Zα (K169A, K170A, R174A) or acidic residues on the poly-E motif should abrogate binding in a predictable manner. Second, cellular co-localization studies and proximity ligation assays could determine whether the interaction occurs in a specific nuclear compartment or under particular transcriptional conditions. Third, it will be important to establish whether YTHDC1 binding is competitive with, cooperative with, or independent of Z-RNA binding at the Zα surface, and whether the interaction regulates ADAR1 editing activity on specific transcripts. It will also be of interest to determine whether acidic patches, such as those on MBD3 and YTDC1, help localize Z-DNA-binding proteins to other DNA and RNA modification complexes.

In summary, we present a computational pipeline that integrates generative protein design, proteome-wide structural screening, and microsecond molecular dynamics to discover that YTHDC1 engages the ADAR1 Zα domain through an acidic protein mimic of Z-RNA. This finding provides a structural basis for cross-talk between innate immunity editing and epitranscriptomic m6A reading, and demonstrates the power of inverse design strategies for unbiased discovery of protein–protein interactions at structurally defined surfaces.

## Acknowledgements

We would like to thank A.S. Fedulova and A.K. Gribkova for help with MDS and valuable discussions of MDS results and AlphaFold3 predictions that helped to improve this work.

## Data availability

All synthetic binder sequences generated by RFdiffusion and ProteinMPNN, ColabFold and AlphaFold3 predicted structures and model confidence files, molecular dynamics input parameters, and custom analysis scripts used in this study are deposited at https://doi.org/10.5281/zenodo.21229987. Representative MD trajectory frames and topology files are included in the Zenodo repository. Full microsecond MD trajectories are available from the corresponding author upon reasonable request. RFdiffusion, ProteinMPNN, ColabFold, and AlphaFold3 are open-source tools available at their respective repositories cited in the Methods section.

